# Competence Inhibition by the XrpA Peptide Encoded Within the *comX* Gene of *Streptococcus mutans*

**DOI:** 10.1101/265223

**Authors:** Justin Kaspar, Robert C. Shields, Robert A. Burne

**Affiliations:** Department of Oral Biology, University of Florida, Gainesville, Florida 32610

**Author notes:** Corresponding author: Mailing address: Department of Oral Biology, University of Florida, College of Dentistry, P.O. Box 100424, Gainesville, FL 32610.

**Keywords:** quorum sensing, intercellular signaling, transcription factor, genetic competence, dental caries

## Abstract

*Streptococcus mutans* displays complex regulation of natural genetic competence. Competence development in *S. mutans* is controlled by a peptide derived from ComS (XIP); which along with the cytosolic regulator ComR controls the expression of the alternative sigma factor *comX*, the master regulator of competence development. Recently, a gene embedded within the coding region of *comX* was discovered and designated *xrpA* (com<u>X</u> *r*egulatory <u>p</u>eptide <u>A</u>). XrpA was found to be an antagonist of ComX, but the mechanism was not established. In this study, we reveal through both genomic and proteomic techniques that XrpA is the first describe negative regulator of ComRS systems in streptococci. Transcriptomic and promoter activity assays in the Δ*xrpA* strain revealed an up-regulation of genes controlled by both the ComR- and ComX-regulons. An *in vivo* protein crosslinking and *in vitro* fluorescent polarization assays confirmed that the N-terminal region of XrpA were found to be sufficient in inhibiting ComR-XIP complex binding to ECom-box located within the *comX* promoter. This inhibitory activity was sufficient for decreases in P*comX* activity, transformability and ComX accumulation. XrpA serving as a modulator of ComRS activity ultimately results in changes to subpopulation behaviors and cell fate during competence activation.

**ABBREVIATED SUMMARY:** *Streptococcus mutans* displays complex regulation of natural genetic competence, highlighted by a novel gene, *xrpA*, embedded within the coding region for the master regulator ComX. We show that XrpA modulates ComRS-dependent activation of *comX* expression, resulting in changes to sub-population behaviors, including cell lysis. XrpA is the first described inhibitor of a ComRS system and, because it is unique to *S. mutans* it may be targetable to prevent diseases caused by this pathogen.

## INTRODUCTION

Bacterial biofilm communities coordinate behaviors in response to environmental stimuli through the use of chemical mediators that accumulate extracellularly to activate transcription of specific genes when a critical concentration is achieved, in a process termed quorum sensing (Fuqua *et al*., 1994). In Gram-negative bacteria, diffusible acylated homoserine lactones are the principal chemical mediator that act as a proxy for cell density (Papenfort and Bassler, 2016), whereas small hydrophobic peptides fulfill a similar role in Gram-positive bacteria (Håvarstein *et al*., 1995). More recently, quorum sensing and gene products involved in intercellular signaling have been highlighted as an area of interest for therapeutic intervention in some bacterial infections, because quorum sensing often controls the transcription of genes that contribute to virulence (Greenberg, 2003).

Genetic competence, a transient physiological state in which bacteria produce the gene products necessary for uptake of DNA from their environment was one of the first described and studied quorum sensing pathways (Tomasz, 1965). Decades later, genetic competence is still a valuable model system in molecular microbiology to unravel the complexities of signal perception, signal transduction and sub-population behaviors, and is of relevance to newer areas of research that include sociomicrobiology, interspecies antagonism and cooperativity, and microbial biogeography (Whiteley *et al*., 2017). Genetic competence has been extensively studied in the genus *Streptococcus*, including *Streptococcus pneumoniae* (Hui *et al*., 1995; Straume *et al*., 2015), *Streptococcus thermophilus* (Fontaine *et al*., 2009; Gardan *et al*., 2013), *Streptococcus pyogenes* (Mashburn-Warren *et al*., 2012; Wilkening *et al*., 2015) and *Streptococcus mutans* (Li *et al*., 2001; Son *et al*., 2012). Interestingly, streptococci harbor two distinct peptide signaling systems that activate genetic competence: the Mitis and Anginosus groups utilize an extracellular signaling system composed of a signaling peptide termed CSP (competence stimulating peptide) and a two-component signal transduction system encoded by *comDE*. In contrast, the Bovis, Salivarius and Pyogenic streptococci employ an intercellular signaling system that consists of the signal peptide XIP (*comX/sigX* inducing peptide) derived from the ComS precursor, and a cytosolic Rgg-like regulator designated as ComR (Håvarstein, 2010). While these two signal systems diverge substantially in their distribution among species and how the systems perceive and transduce their signals, stimulation of either pathway with the cognate peptide results in the activation of transcription of an alternative sigma factor termed ComX or SigX. ComX controls the transition into the competent state by activating the expression of a regulon encoding gene products necessary for DNA uptake and processing.

The Mutans group of streptococci, including the human caries pathogen *S. mutans*, are unique in that most strains encode an apparently functional ComCDE as well as ComRS pathways. Further, either signal peptide (CSP or XIP) can trigger up-regulation of *comX*, although different conditions, including pH, redox and growth phase, influence how effectively each pathway is able to function (Hagen and Son, 2017). Addition of synthetic CSP (sCSP) to growing cultures of *S. mutans* in a peptide-rich medium, such as BHI, results in activation of transcription of genes for the biogenesis of bacteriocins via direct binding of phosphorylated ComE to a conserved sequence in the promoter regions of these genes and operons. Consistent with this observation, the ComCDE system of *S. mutans* appears to have evolved from a common ancestor of the BlpCRH system of S. *pneumoniae*, which does not regulate competence but does induce bacteriocins in the pneumococcus and some related organisms (Johnston *et al*., 2014). Transcription of *comX* can also be induced by CSP, but this generally occurs in only a subset of organisms in a population, it does not involve direct binding of ComE to the *comX* promoter, and the underlying mechanism for CSP-dependent activation of *comX* is not well-understood (Kreth *et al*., 2007; Hung *et al*., 2011). The proximal regulator for direct activation of competence is ComRS. When synthetic XIP (sXIP) is added to a peptide-free, chemically defined medium, such as FMC or CDM, it is imported into the cytosol by the oligopeptide permease OppA and forms a complex with ComR to activate *comX* transcription in the entire bacterial population (Mashburn-Warren *et al*., 2010; Son *et al*., 2012). The XIP-ComR complex can also activate the gene for the precursor of XIP, *comS*, creating a positive feedback loop for amplification of the competence activation signal (Fontaine *et al*., 2013). XIP has been detected in culture supernates, supporting the hypothesis that XIP is a diffusible intercellular signal (Desai *et al*., 2012; Khan *et al*., 2012; Wenderska *et al*., 2012), although the mechanism by which XIP is released into the environment can involve active transport (Chang and Federle, 2016) or cell lysis (Kaspar *et al*., 2017), depending on the species of bacteria. Recently, it was confirmed experimentally that XIP is able to act as a diffusible intercellular communication molecule and that signaling can occur within biofilm populations (Shields and Burne, 2016; Kaspar *et al*., 2017).

While substantial progress has been made dissecting the mechanisms leading to *com* gene activation, very little is known about regulation of the system after *comX* is induced and late competence gene expression is active. In S. *pneumoniae*, shut-off of the ComCDE system is regulated at multiple levels, including competition between phosphorylated and un-phosphorylated ComE for binding sites, direct inhibition of activated ComE by the late competence gene-encoded protein DprA, and by inhibition of ComX activity by an unknown factor (Martin *et al*., 2013; Mirouze *et al*., 2013; Weng *et al*., 2013). In streptococci that harbor ComRS systems, the factors that regulate the Com circuit after transcriptional activation have not been characterized in significant detail. It has been postulated that a “*comZ* gene”, under the control of ComX, exists that encodes a product that acts on the ComR-XIP complex to create a feedback inhibition loop (Boutry *et al*., 2013; Haustenne *et al*., 2015). Recently, we described a novel protein encoded within *comX* gene in an alternative (+1) reading frame that we designated as XrpA *(comX* regulatory protein/peptide A) (Kaspar *et al*., 2015). We described XrpA as a novel antagonist of *comX* as loss of XrpA, either by mutating the start codon or by introducing premature stop codons, led to an increase in transformation efficiency and accumulation of the ComX protein, whereas overexpression of *xrpA* resulted in decreased transformability and lower levels of ComX. However, the mechanism by which XrpA exerted its effects was not determined. In this study, we present genetic evidence through transcriptome profiling and biochemical evidence of protein-protein interactions that demonstrate that XrpA affects competence development in *S. mutans* by interacting with and inhibiting ComR activity, thus describing the first negative regulator of competence signaling that acts on the ComRS circuit.

## RESULTS

### Transcriptome profiling of an XrpA-deficient strain

In our initial characterization of *xrpA*, we highlighted the unusual transcriptional characteristics of *xrpA* and the profound influence of XrpA on the dramatically different genetic competence phenotypes displayed by strains with polar and non-polar mutations in the *rcrR* gene of the *rcrRPQ* operon, designated Δ*rcrR*-P and Δ*rcrR*-NP, respectively. Inactivation of *xrpA* in a way that did not alter the primary sequence of ComX could convert the non-transformable Δ*rcrR*-NP strain into the hyper-transformable state that was observed for the ΔrcrR-P strain, with concomitant restoration of ComX production (Kaspar *et al*., 2015). However, the *rcrR* mutants displayed extreme and unusual phenotypes, so questions remain as to how *xrpA* expression is regulated and what role XrpA plays in a wild-type *S. mutans* genetic background. To begin to answer these questions, we first wanted to compare the transcriptomes of a Δ*xrpA* strain with that of the *S. mutans* wild-type strain, UA159. For these studies and those conducted henceforth, the Δ*xrpA* used contains a single base change at the 162^nd^ nucleotide of *comX* (*comX*::T162C), which mutates the *xrpA* start codon (ATG→ACG) and leaves the *comX* protein coding sequence unchanged (Kaspar *et al*., 2015).

Comparison of the transcriptome by RNA-Seq of the wild-type with the mutant lacking XrpA when cells were grown in the chemically defined medium FMC to mid-exponential phase revealed 56 differentially expressed genes, with 34 upregulated in Δ*xrpA* and 22 downregulated (**Figure 1A**). Many of the upregulated genes were competence-related genes, including *comX* and genes that are a part of the ComX regulon of *S. mutans* (Khan *et al*., 2016) (**Table S2**). Since loss of *xrpA* caused upregulation of competence genes, we also analyzed the transcriptome of UA159 treated with 2 μM sXIP to induce competence, with UA159 treated with vehicle (DMSO) as a control. Cells treated with sXIP had 137 genes differentially expressed compared to the control (**Figure 1B; Table S3**). Several of the same genes that were the most strongly upregulated in Δ*xrpA*, including the *comF* and *comY* operons, *drpA* and *lytF*, were also the highest upregulated genes in the sXIP-treated cells. Interestingly, those genes that were downregulated in the Δ*xrpA* strain differed from those downregulated by sXIP addition to UA159. All of the downregulated genes in the Δ*xrpA* mutant were located on the TnSMu1 genomic island that encodes predicted transposases, integrases, transporter(s) and hypothetical proteins (Waterhouse *et al*., 2007). This genomic island was recently found to be differentially expressed in *clpP* and *cidB* mutants, providing additional evidence for a link between XrpA and stress responses (Chattoraj *et al*., 2010; Kaspar *et al*., 2015; Ahn and Rice, 2016).

**Figure 1.**
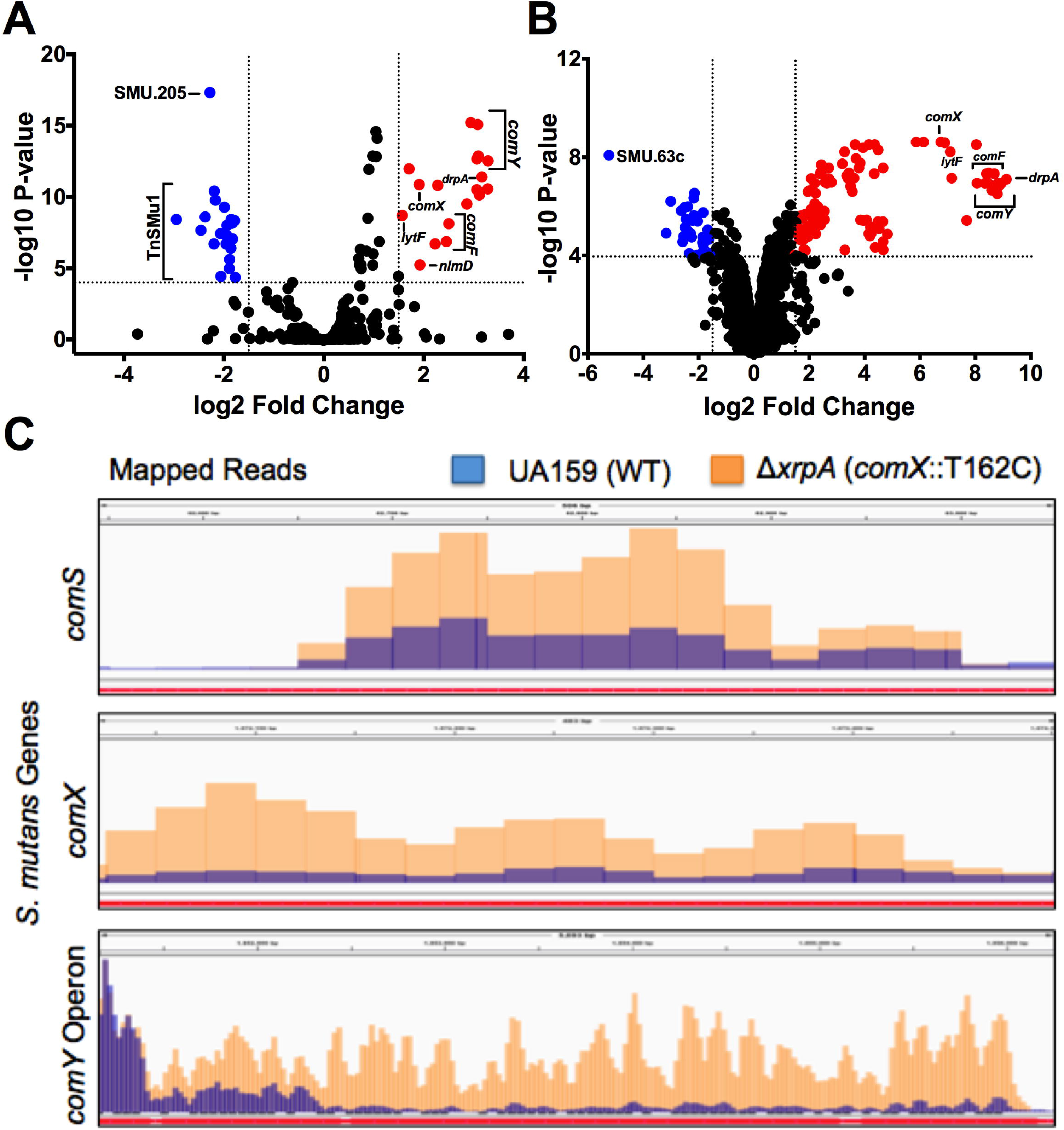
*Transcriptome Analysis of ΔxrpA and UA159 with sXIP addition*. Volcano plots of (A) Δ*xrpA* and (B) UA159 + 2 μM sXIP from RNA-Seq results compared to UA159 of three independent replicates grown in FMC medium to OD_600_ = 0.5. Log2 fold change and false discovery rates (FDR) converted to −log10 P-values were calculated from Degust using edgeR analysis. Genes of interest that were ≥1.5 log2 fold change and had a ≥4 −log10 P-value were highlighted either in red (upregulated) or blue (downregulated) and are listed in Tables S2 and S3. (C) Visual representation of read counts accumulated in either the *comS, comX* or *comY* coding sequences in either the UA159 (blue bars) or Δ*xrpA* (orange bars) genetic backgrounds. Mapped short read alignments were converted in “.bam” files and visualized with the IGV genome browser.

Meanwhile, the upregulation of the ComX regulon in the Δ*xrpA* mutant could not be explained by increases in expression of annotated early competence genes. However, when we examined the region encoding *comS*, which is encoded in the intergenic region of SMU.61 and SMU.63c in current database annotations, we found an elevated number of reads for *comS* in the Δ*xrpA* strain compared to UA159 (**Figure 1C**). When one considers that both *comS* and *comX* are upregulated in the XrpA-deficient strain, a clearer picture emerges that XrpA most likely exerts its influence over competence development by influencing the efficiency of ComR-dependent activation of the *comX* and *comS* promoters, P*comX* and P*comS*.

### XrpA alters comX and comS promoter activity

To determine if XrpA affects the ComR-XIP activated promoters, PcomX and PcomS, we incorporated the *xrpA* start codon mutation *comX*::T162C into strains carrying GFP transcriptional fusions to each promoter. Similarly, we transformed an xrpA-overexpressing strain (184XrpA) into the GFP reporter gene fusion strains to see if increasing the amount of XrpA produced would yield gene expression patterns that were opposite of those caused by loss of *xrpA*. Cells were grown in chemically defined CDM, which allows for self-activation of the ComRS system. Indeed, loss of *xrpA* resulted in both earlier and higher level expression of P*comX* and *PcomS* reporters, compared to what was observed in the wild-type genetic background. Conversely, overexpression of *xrpA* from the strong, constitutive promoter on pIB184 caused a decrease in GFP production from the *comX* and *comS* promoters (**Figure 2**). Collectively, these results validate the RNA-Seq data and provide support for the hypothesis that XrpA negatively affects the activation of gene expression by ComR.

**Figure 2.**
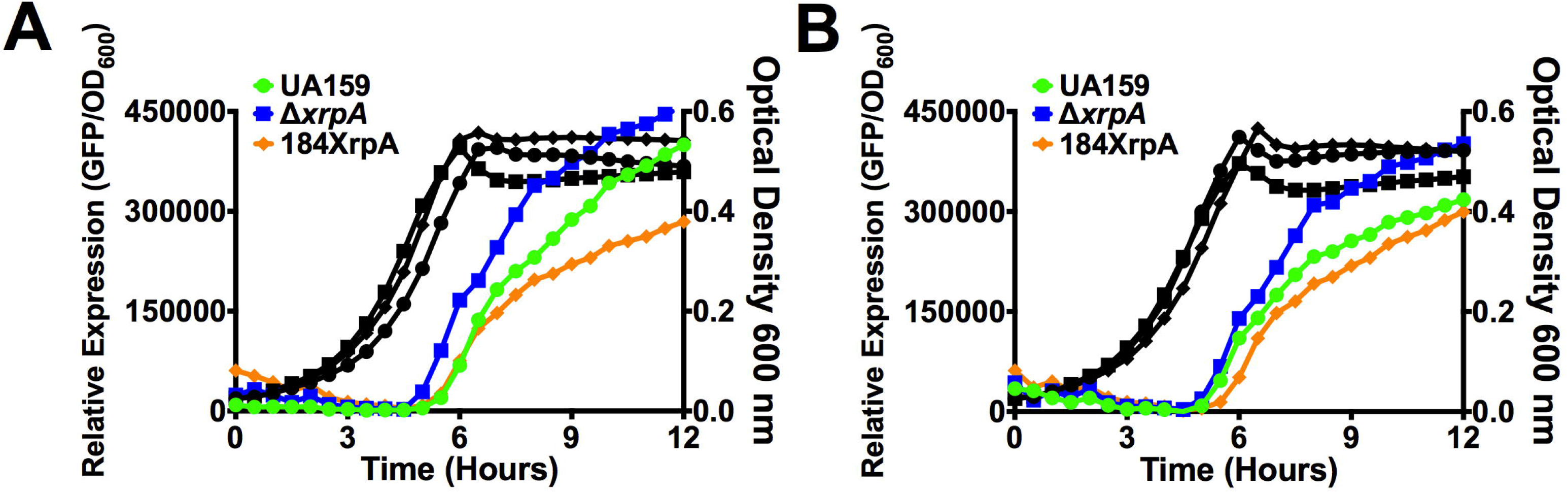
*Effect of XrpA on activation of* comX *and* comS *promoters*. Transcriptional activation assays using a fused-gfp reporter for (A) P*comX* and (B) *PcomS* in wild-type (UA159 – green circles), comX::T162C *(ΔxrpA* – blue squares) or *xrpA* overexpression (184XrpA – orange diamonds) genetic backgrounds. Black lines represent growth (OD_600_, right axis) of each of the respective reporter strains during the assay. Each data point shown is the average of three independent biological replicates with four measured technical replicates.

### XrpA influences subpopulation responses to competence signals

Stimulation of genetic competence in a peptide-rich medium, such as brain-heart infusion (BHI), by CSP results in a bimodal response where only a sub-population of cells activate P*comX* (Son *et al*., 2012). We reasoned that XrpA might influence the proportion of cells that responded to sCSP in BHI. We utilized both the wild-type and Δ*xrpA* mutant carrying P*comX* transcriptional reporter gene fusions and analyzed subpopulation behaviors using flow cytometry three hours after sCSP addition to planktonic cultures. The percentage of cells that were GFP-positive was more than 20% greater in the Δ*xrpA* background with addition of 100 nM sCSP (39.7 ± 2.2 versus 61.5 ± 1.8) (**Figure 3A**) or 1000 nM sCSP (48.7 ± 1.4 compared to 73.9 ± 2.3) (**Figure 3B**). Further, mean GFP fluorescence intensity was increased in the Δ*xrpA* background.

**Figure 3.**
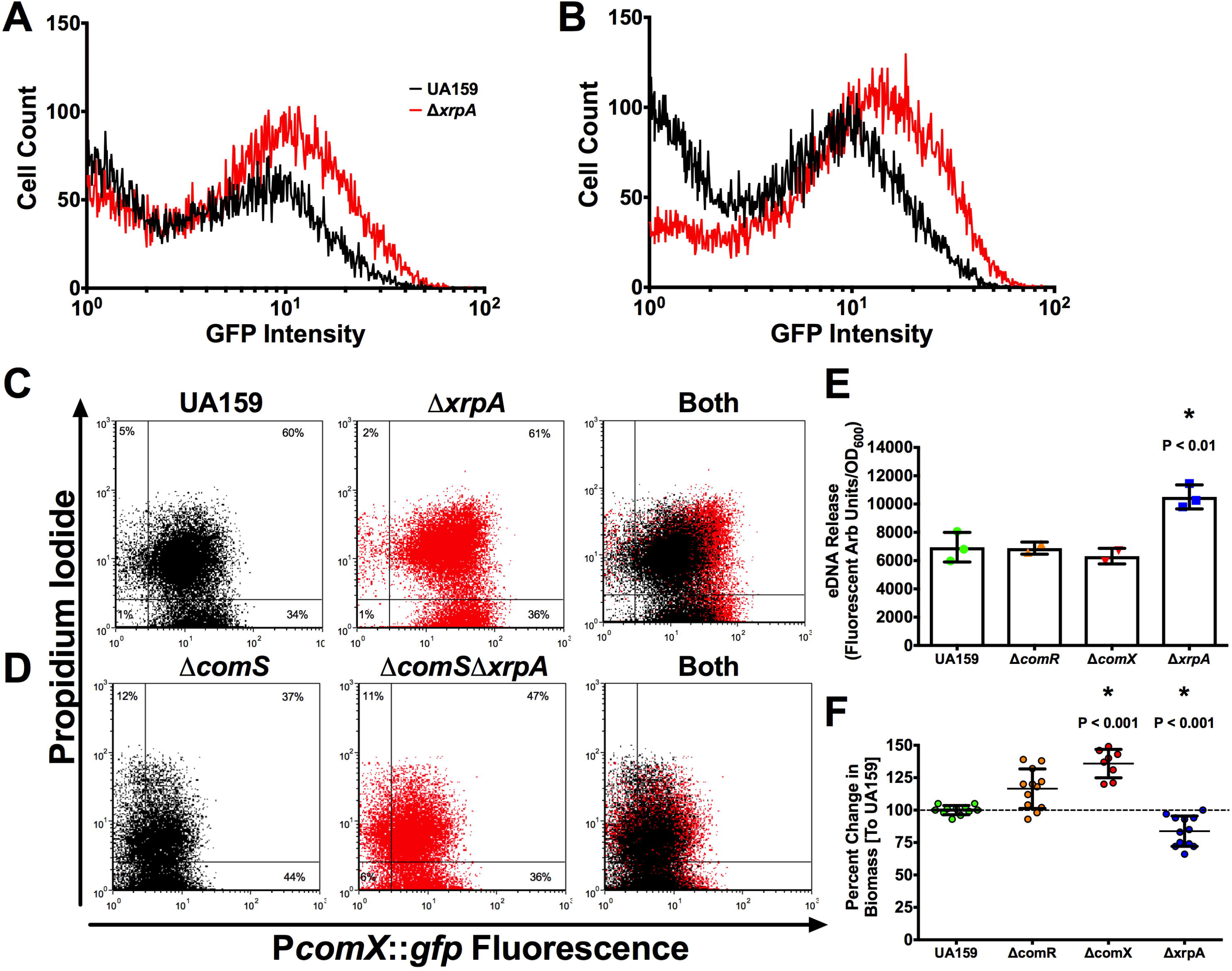
*XrpA changes subpopulation behaviors*. Histogram of cell counts from flow cytometry analysis of the *PcomX::gfp* reporter strain in UA159 (black lines) or Δ*xrpA* (red lines) grown in BHI with addition of either (A) 100 nM or (B) 1000 nM sCSP. For experiments with FMC and addition of 2 μM sXIP, reporter strains in either (C) UA159 or (D) Δ*comS* background were stained with propidium iodide before analysis. A total of 50,000 cells were counted in three independent replicates for each experiment. (E) eDNA release of selected strains from three independent overnight cultures grown in CDM media. eDNA release was calculated by taking the arbitrary fluorescent units and dividing by the recorded OD_600_ at the time of harvest. (F) Change in biofilm biomass compared to UA159 using CDM media with 15 mM glucose and 2.5 mM sucrose as the carbohydrate source after 48 hours of growth. Each data point (n = 12 or n = 8) represents an individual replicate. Statistical analysis was calculated by the Student’s *t*-test; * P < 0.01.

Stimulation of the competence cascade by XIP in nanomolar concentrations of wild-type *S. mutans* growing in a peptide-free medium results in a unimodal population response, but addition of sXIP to cultures of *S. mutans* at concentrations higher than 1 μM can trigger cell death in a significant fraction of the population (Wenderska *et al*., 2012). To determine if XrpA could influence XIP-mediated killing in cells treated with higher concentrations of sXIP, we followed a protocol similar to the CSP experiments, but stained the cells with propidium iodide (PI) to measure membrane integrity prior to analysis by flow cytometry. No changes were seen in the proportions of the population that were PI-positive between the wild-type and Δ*xrpA* background, but a clear increase in the mean GFP intensity was observed when *xrpA* was mutated, similar to what was seen with CSP (**Figure 3C**). However, when the *comS* gene was removed to eliminate the positive feedback loop in the XIP signaling pathway, a distinct increase in the proportion of PI-positive cells was seen in the Δ*xrpA* mutant population, compared to behaviors in the wild-type genetic background (36.8 ± 0.7 versus 47.2 ± 0.9) (**Figure 3D**). Measurements of eDNA release from overnight cultures were used to confirm the finding that strains that were activated for competence, but that lack *xrpA*, were more lytic than their wild-type counterparts (**Figure 3E**). This lytic behavior could also be measured in a biofilm model, where the absence of XrpA correlated with decreased biofilm formation after 48 h, compared to increased biofilm formation in the non-competent Δ*comR* or Δ*comX* mutant strains (**Figure 3F**). Taken together, these data highlight that XrpA can influence subpopulation behaviors both in terms of competence activation in environments where peptides are present, and influence lytic behaviors associated with activation of the competence by high, albeit physiologically relevant, concentrations of signal peptide, with the enhanced cell death most likely being associated with more robust activation of P*comX*.

### Addition of XrpA can inhibit competence development

As data from transcriptome profiling and transcriptional reporter experiments support a role for XrpA in interference with the ComRS pathway, we wanted to test whether direct addition of XrpA could inhibit competence development. We tried various strategies to produce a full-length recombinant XrpA in *Escherichia coli*, however we were unsuccessful, possibly due to the combination of its hydrophobicity and having 7 cysteines distributed across the protein. As a substitute, we settled on synthesizing four different peptide fragments that span the entirety of the 69-aa XrpA protein (**Figure 4A**). We first tested the ability of the peptides to interfere with activation of the P*comX* transcriptional reporter in cells growing in CDM. Of the four peptides tested, only XrpA-1 containing aa 5-20 of the N-terminal region of the protein caused a marked decreased in P*comX* activity, compared to when only vehicle was added (**Figure 4B**). Interestingly, a higher final OD_600_ was also recorded for cultures exposed to XrpA-1, compared to when either vehicle or any of the other three peptides were tested. To confirm that this effect was specific for XrpA-1, a scrambled peptide (XrpA-S1; same aa composition, different sequence) was synthesized and tested (**Figure 4A**). While some residual inhibitory activity was evident with the scrambled peptide, compared with control and other peptides, the effects of the scrambled peptide was much less pronounced than XrpA-1 (**Figure 4C**). The inhibitory effect of the XrpA-1 peptide was also found to be dose-dependent in this experimental setup (**Figure 4D**). A similar profile for XrpA-1 was seen using the ComR-XIP activated P*comS::gfp* transcriptional fusion strain (data not shown). Together, these results confirm that selected XrpA fragments can inhibit ComRS-dependent activation of *comX* when provided exogenously to cells growing in a chemically defined medium. It is important to note that we presently have no data to support that XrpA peptides are excreted or follow a similar lifecycle as XIP. We contend that observing a response by providing the peptides exogenously was a fortuitous occurrence and future studies will be needed to understand the mechanism for the effect.

**Figure 4.**
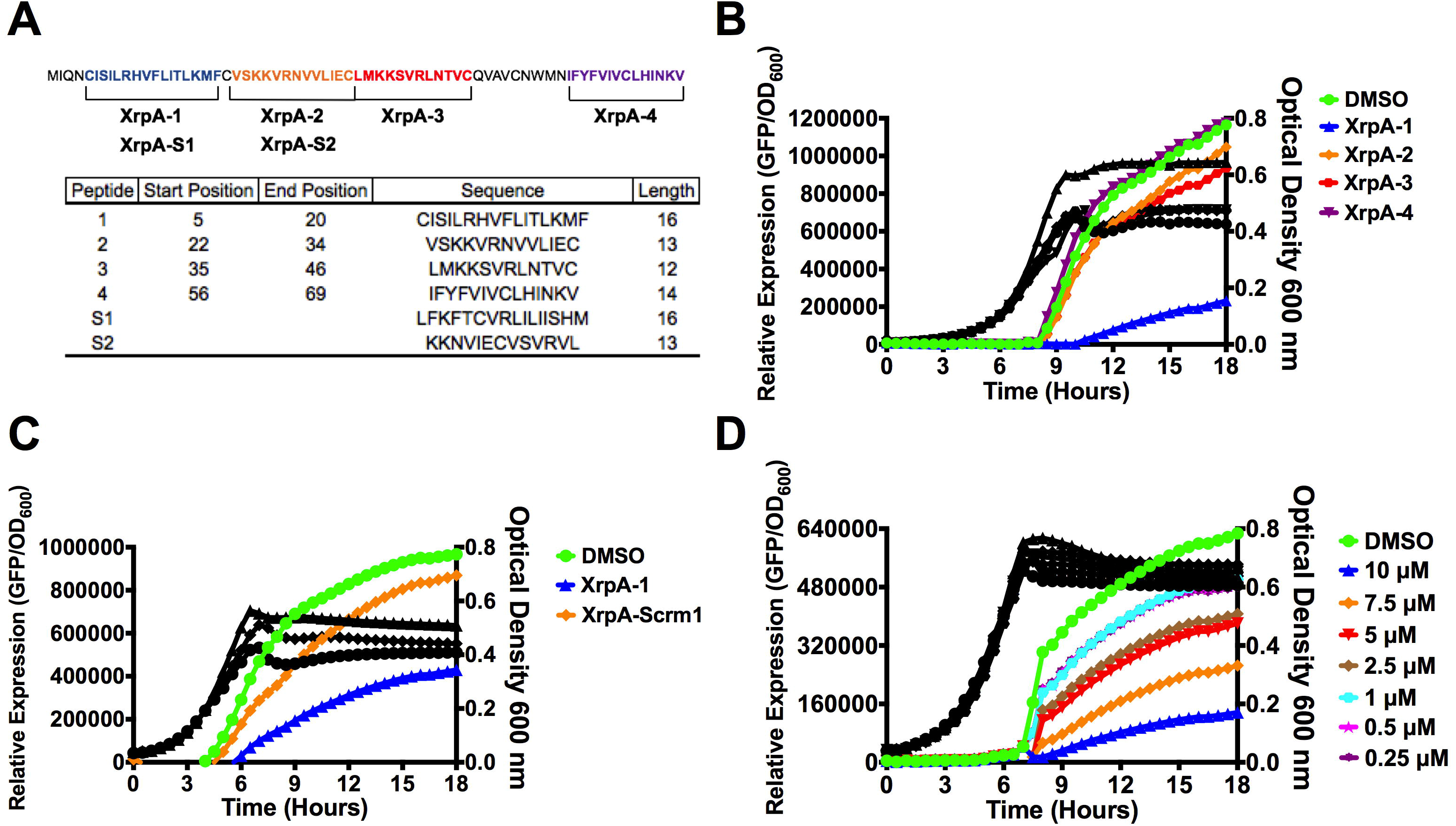
*PcomX activities with addition of various synthetic XrpA peptides*. (A) Synthetic peptides synthesized for use in this study. The full-length, 69-aa XrpA protein is shown with each selected peptide highlighted by color and brackets. Below is a table with each peptide start and stop position in native XrpA, their sequences, and length in aa residues. Peptides S1 and S2, representing scrambled XrpA-1 and XrpA-2, were generated by the Shuffle Protein tool from www.bioinformatics.org with XrpA-1 and XrpA-2 serving as input sequences; S1 and S2 are of same length and aa composition as XrpA-1 and -2, respectively. (B) Transcriptional activation assays using *PcomX*::*gfp* reporter strain grown in CDM medium with addition of 10 μM of various synthesized XrpA peptides. (C) Direct comparison of transcriptional activation assays between XrpA-1 and a scrambled version of XpA-1. (D) Dose-dependent inhibition of P*comX* activity by various concentrations of XrpA-1. Colored lines represent relative P*comX* expression (arbitrary fluorescence units divided by OD_600_, left axis). Black lines represent growth (OD_600_, right axis) of each of the respective reporter strains during the assay. Each data point shown is the average of three independent biological replicates with four measured technical replicates.

An explanation for the higher OD_600_ values when XrpA-1 was added to the growing cultures may be related to reduced cell lysis from decreased competence activation. eDNA release was measured as described for **Figure 3 and we found significantly less eDNA accumulation in culture supernates from overnight cultures grown in the presence of XrpA-1, but not with the other XrpA peptides, compared to DMSO-treated controls. (Figure 5A**). We also tested several other competence-related phenotypes with the synthetic XrpA peptides. In terms of transformation efficiency, a significant −1.22 ± 0.05-Log10 fold decrease was observed when XrpA-1 was present, compared to the DMSO only control, and −0.23 ± 0.07 and −0.08 ± 0.19-Log_10_-fold decreases were observed when XrpA-2 and XrpA-3 were present, respectively (**Figure 5B**). In terms of ComX accumulation, ComX protein was not observed by western blotting in cells treated with XrpA-1 (**Figure 5C**). It did not appear as if ComR levels were impacted by addition of any of the peptides, suggesting that XrpA does not influence ComRS activity by altering ComR stability. In all, these results support the gene fusion assays and show that XrpA addition to growing cultures has a significant impact on competence development and lysis-related phenotypes when provided exogenously to *S. mutans*.

**Figure 5.**
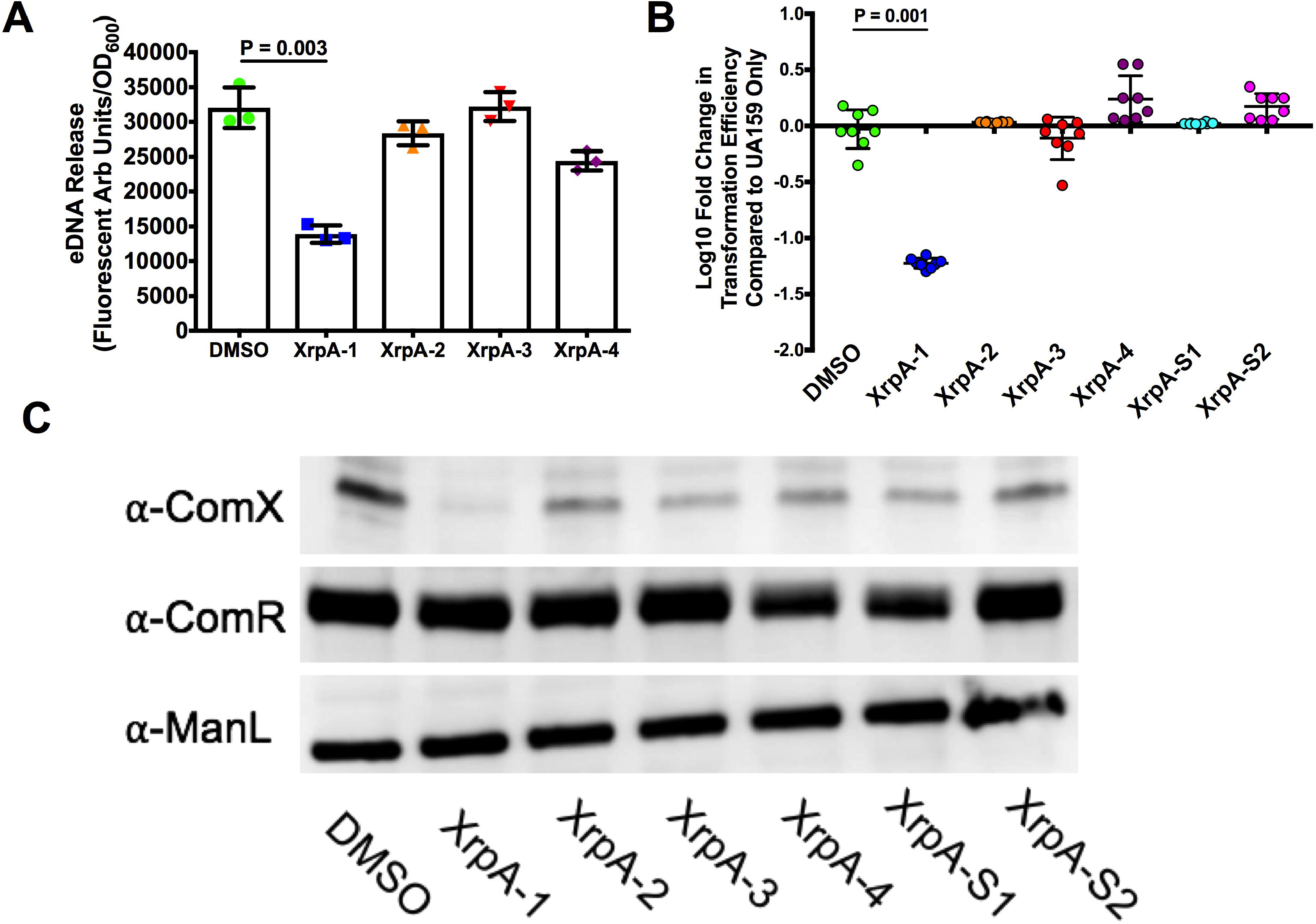
*Addition of XrpA peptides changes lytic and competence phenotypes*. (A) eDNA release by UA159 in cultures to which 10 μM XrpA peptides were added. Results are from three independent overnight cultures grown in CDM media. eDNA release was calculated by taking the arbitrary fluorescence units and dividing by the recorded OD_600_ at the time of harvest. (B) Log_10_ fold change in transformation efficiency with 10 μM XrpA peptides compared to DMSO-only control (vehicle; set to 0) in FMC medium with 0.5 μM sXIP addition. Each data point represents a replicate (N = 8). (C) Western blot using 10 μg of whole cell lysates of UA159 with addition of 10 μM XrpA peptides in FMC medium with 0.5 μM sXIP addition. Cells were grown to OD_600_ = 0.5 before harvesting. Primary antisera, raised against the corresponding protein, were used for detection. ManL (the EIIAB domain of the glucose PTS permease) served as a loading control. All statistical analysis for this figure was calculated by the Student’s *t*-test.

### In vitro interactions of XrpA and ComR

We hypothesized that XrpA inhibits competence development through direct interaction with ComR, affecting ComR DNA binding activity and leading to reduced P*comX* activation. To confirm this hypothesis, we utilized a fluorescence polarization (FP) assay where we could monitor the binding of the ComR-XIP complex to the promoter region of *comX in vitro* using purified ComR protein and a 5’ Bodipy-labeled, self-annealing stem-loop DNA probe that encompassed the ECom-box to which ComR-XIP binds for transcriptional activation (Mashburn-Warren *et al*., 2010; Fontaine *et al*., 2013). Strong, direct binding of ComR-XIP was observed to the probe in the absence of any of the XrpA peptides, with a calculated *K_d_* of 153 ± 10 nM (**Figure 6A, Table 2**). However, when XrpA-1 was added to the reaction, the *K_d_* increased to 651 ± 99 nM. Surprisingly, XrpA-2, which had no observable effects on the phenotypes examined above, also displayed inhibitory effects, while inclusion of XrpA-3 or XrpA-4 did not substantially alter the calculated *Kd* values (Table 2). Two different scrambled peptides as detailed in **Figure 4A**, containing the same aa composition as their respective counterpart but different sequence, were tested in combination with XrpA-1 and XrpA-2 to verify the specificity of the effect. Similar to the transcriptional reporter assays, XrpA-S1 showed some alleviation of inhibitory properties (Figure 6B), whereas inclusion of the XrpA-2 scrambled peptide yielded a ComR-XIP DNA binding affinity similar to the control (Figure 6C, Table 2). Taken together, these experiments verify that XrpA interacts directly with ComR, with the outcome being that this interaction antagonizes promoter binding and ultimately gene expression by the ComR-XIP complex and competence development.

**Figure 6.**
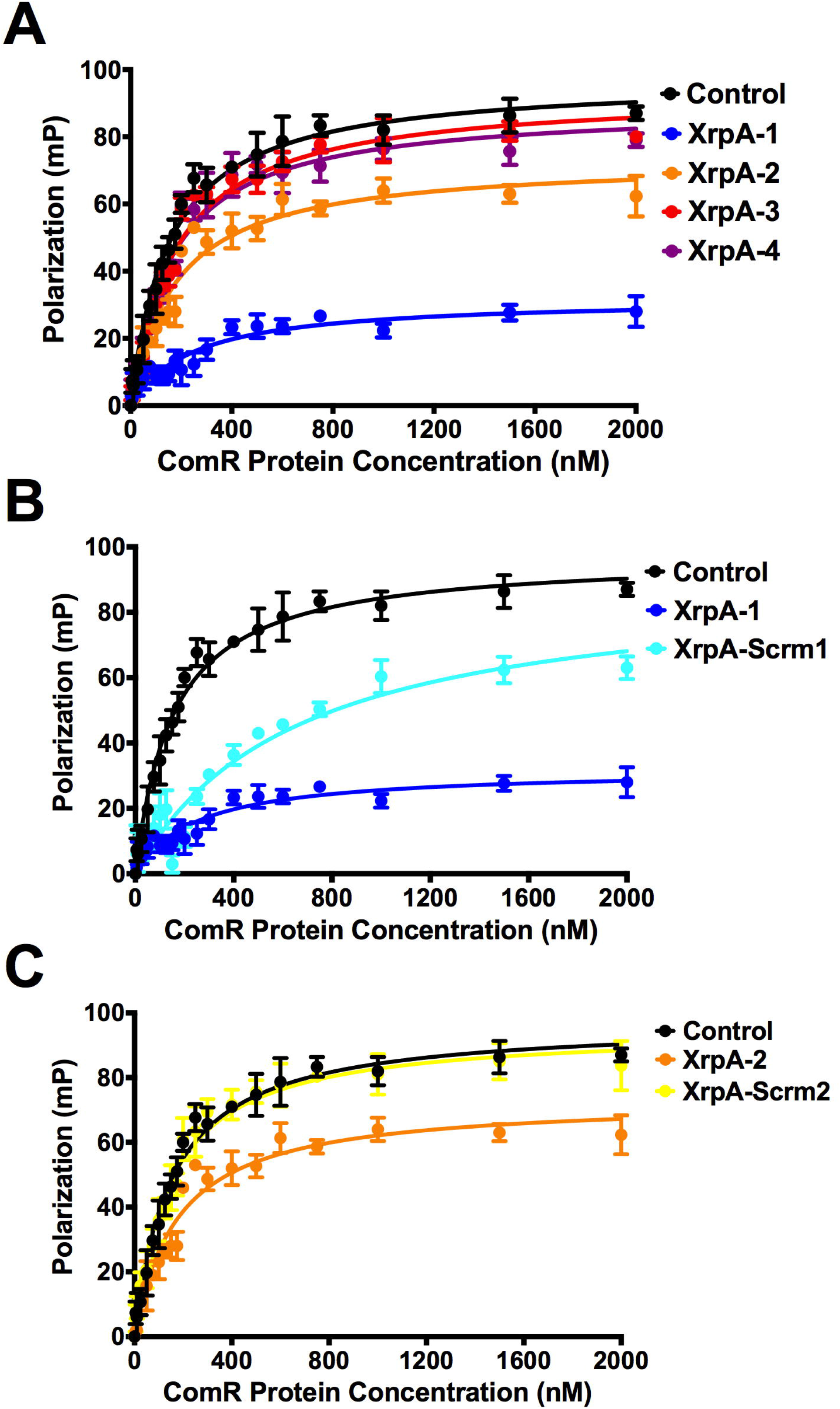
*Fluorescence Polarization confirms XrpA-ComR interactions*. Fluorescence polarization (FP) curves of increasing concentrations of purified ComR binding to 10 nM of P*comX* dsDNA probe in the presence of 10 μM sXIP and 10 μM of various XrpA peptides (Figure 4A). Control (black lines) represent binding in the absence of XrpA peptides. (A) addition of XrpA peptides 1-4, (B) comparison between XrpA-1 and a scrambled version of XrpA-1, and (C) comparison between XrpA-2 and a scrambled version of XrpA-2. Data shown are averages from three independent experiments. *K_d_* values are shown in Table 2.

### The N-terminal region of XrpA directly interacts with ComR

Using our four synthesized peptide fragments of XrpA, it appeared that the N-terminal region of XrpA was the domain responsible for the inhibitory activity. To confirm that the *in vitro* protein-protein interaction data accurately represented *in vivo* activity, we adapted a Strep-protein interaction (SPINE) protocol (Herzberg *et al*., 2007) that allows for crosslinking of proteins *in vivo*, followed by affinity purification of target protein complexes and identification of interacting partners using mass spectrometry (MS). For this experiment, we chose ComR as the bait by incorporating a C-terminal Strep-tag^®^ in front of the stop codon. The construct was also engineered in such a way as to introduce 9 amino acids to serve as a flexible linker sequence to minimize the potential for disrupting the native conformation of ComR; interaction of small hydrophobic peptides by Rgg-like regulators occurs with the C-terminal domain of the proteins (Talagas *et al*., 2016). Cultures (500 ml) of a strain carrying the Strep-tagged ComR expressed from the strong constitutive promoter on pIB184, and a vector-only control strain, were grown in CDM to mid-exponential phase (OD_600_ = 0.6), at which point either 2 μM sXIP or an equivalent volume of 0.1% DMSO control were added. After growth for an additional hour (OD_600_ = 0.8), the homobifunctional *N*-hydroxysuccimide ester (DSP) cross-linking agent that is primary amine-reactive and contains a thiol-cleavable bridge was added to the cells for 45 minutes at 37°C, cells were harvested, and clarified whole cell lysate were passed over a Strep-Tactin^®^ resin for isolation of the targeted complex, as detailed in the methods section (**Supplemental Figure 1**).

In our initial experiments, purified protein complexes were subjected to SDS-PAGE, followed by silver staining (**Supplemental Figure 2**). Five bands of interest that appeared in the Strep-tagged ComR sample, but not in the vector control sample, were excised from the gel and identified by LC-MS/MS after trypsin digestion (**Supplemental Table 5**). Peptide fragments were identified that were derived from the transcriptional regulator SgaR (SMU.289) and a putative single-stranded DNA binding protein Ssb2 encoded by SMU.1967, as well as peptides derived from XrpA. In a second experiment, the purified protein complexes were subjected to two-dimensional differential gel electrophoresis (2D DIGE) (**Figure 7A, Supplemental Figure 3**). A total of 58 spots of interest were selected and a protein expression ratio (PER) was calculated between triplicate samples (**Supplemental Table 4**). Spots chosen for identification were required to have a PER >1.5 compared to the vector-only control sample. The selected spots, 28 in total, were then excised from the gel, digested by trypsin, peptide sequences identified by LC-MS/MS and then compared to a list of predicted masses for all *S. mutans* proteins. While several *S. mutans* proteins were identified with high confidence (**Supplemental Table 5**), peptides derived from XrpA were identified from the 2D DIGE and the fragments detected originated exclusively from the N-terminus of XrpA (aa 1-38), with the fragment MFCVSKK being the peptide that was most frequently observed. Notably, one of the most frequently identified peptide fragments of XrpA observed in the SPINE experiment was not part of the sequence of XrpA-1, so we speculated that using larger peptides might elicit a stronger inhibitory effect. Peptides N1 and N2 were synthesized that included aa 1-18 and aa 18-38, respectively, encompassing all identified XrpA fragments (**Figure 7B**). Indeed, XrpA-N1 substantially decreased P*comX* activity, similar to XrpA-1, whereas addition of XrpA-N2 had an intermediate effect compared to the DMSO control (**Figure 7C**). Similar to the growth profiles of the transcriptional reporter strains seen in **Figure 4B**, XrpA-N1 also displayed better growth, compared to when XrpA-N2 or vehicle was added. These results were mirrored in fluorescent polarization experiments with recombinant ComR and the P*comX* probe (Figure 7D). In all, these experiments implicate the N-terminal region of XrpA in interactions with ComR.

**Figure 7.**
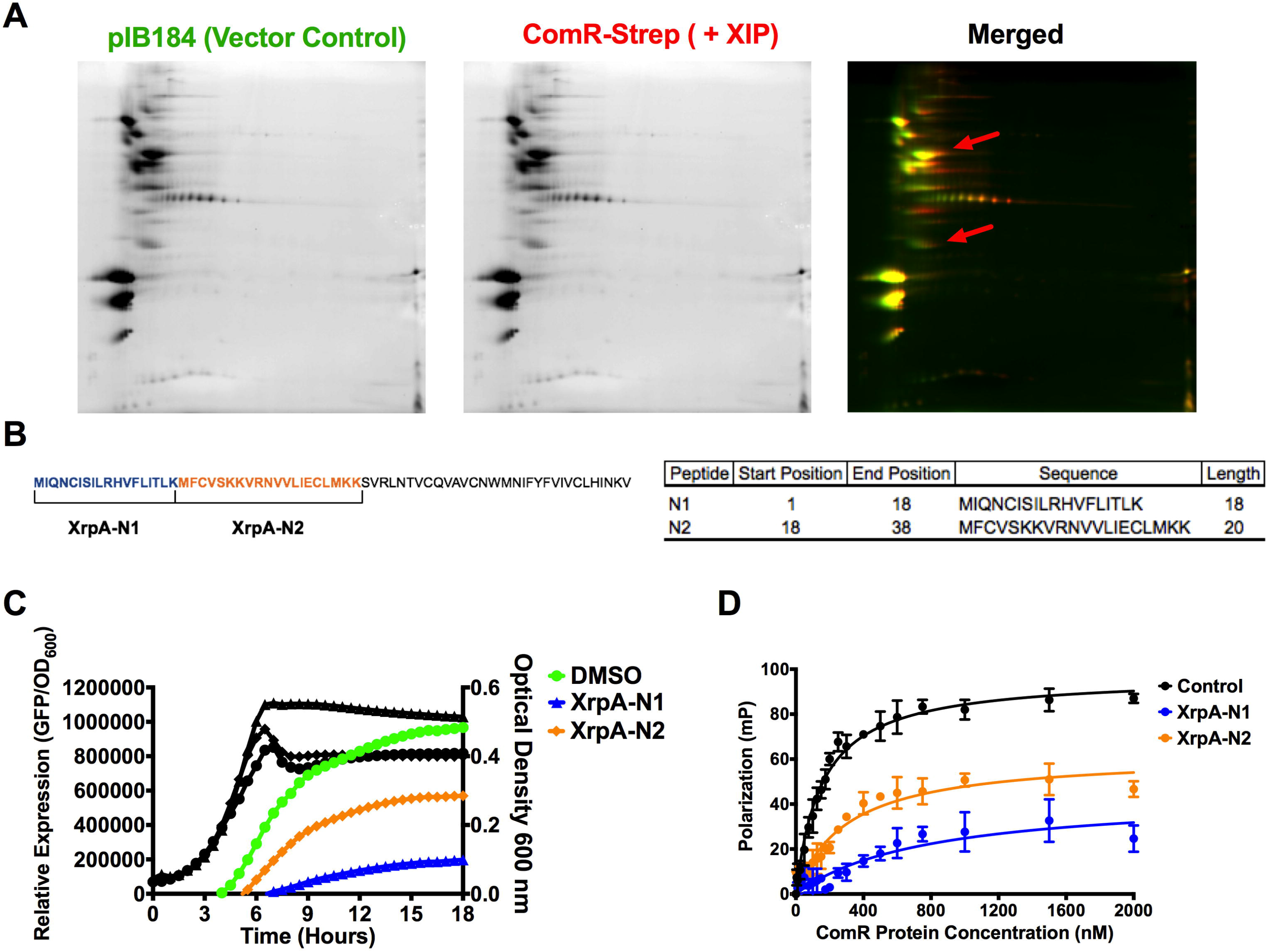
*The N-terminus of XrpA inhibits ComR*. (A) Individual 2D gels of pIB184 (vector control; green) and ComR-Strep with 2 μM sXIP (red) elutions obtained during SPINE experiments, along with a merged image. The red arrows indicate regions where XrpA peptides were identified via spot picking and LC-MS/MS of trypsin-digested proteins. (B) Selection of XrpA-N1 and XrpA-N2 synthetic peptides based on the sequences returned by LC-MS/MS. The full length 69-aa XrpA protein is shown with each selected peptide highlighted by color and brackets. To the right is a table with the start and stop position of each peptide in XrpA, the peptide sequence, and the length in aa residues. (C) Transcriptional activation assays using a fused *PcomX::gfp* reporter strain in CDM medium with addition of 10 μM of either XrpA-N1 or XrpA-N2 compared to DMSO control. (D) Fluorescence polarization (FP) curves of increasing concentrations of purified ComR binding to 10 nM of P*comX* dsDNA probe in the presence of 10 μM sXIP and 10 μM of either XrpA-N1 or XrpA-N2 compared to DMSO control. *K_d_* values are shown in Table 2. Both transcriptional activation assay and FP assay results are averages from three independent experiments.

### Binding of XrpA to ComR is not dependent on the <u>comX</u> promoter

While the results described thus far suggest that N-terminal fragments of XrpA are sufficient to diminish the ability of ComR to bind DNA, the effect could be exerted in multiple ways. For example, (1) XrpA could interact directly with the ComR-XIP oligomer(s) to decrease the affinity of the complex for DNA, (2) XrpA could compete with XIP for the SHP (XIP) binding site, or (3) XrpA may inhibit ComR-dependent activation of gene expression by preventing ComR-XIP from forming higher-order oligomeric complexes; the ComR-XIP complex forms dimers upon XIP binding (Fontaine *et al*., 2013). To further explore how XrpA influences ComR behavior, we synthesized a fluorescein isothiocyanate (FITC)-labeled XrpA-N1 peptide and monitored its direct binding to ComR by FP in the absence of the P*comX* DNA probe. We tested the specificity of the FITC-labeled XrpA-N1 peptide probe interaction with ComR by doing fluorescent polarization experiments with two other *S. mutans* purified proteins that do not participate directly in the regulation of competence, CcpA (Abranches *et al*., 2008) and SppA (SMU.508; Zeng, personal communication), and neither showed any ability to interact with fluorescent XrpA-N1 (**Figure 8A**). We also conducted a cold competition FP assay and found poorer binding for our peptide probe to ComR as increasing concentrations of unlabeled XrpA-N1 peptide were added (**Figure 8B**). In terms of experiments with ComR, fluorescence polarization in the presence of XrpA-N1 was increased regardless of whether sXIP was present or absent (**Figure 8C**), indicating that XrpA may interact directly with ComR and that this interaction can occur even if XIP does not occupy the SHP-binding pocket. Finally, addition of increasing concentrations of sXIP had no effect on fluorescent XrpA-N1 affinity for ComR (**Figure 8D**), suggesting that XrpA and sXIP do not compete for the SHP (XIP) binding site. Collectively, these experiments verify that XrpA interacts directly with ComR independent of both ComR-regulated promoters (such as PcomX) as well as XIP, favoring a model where XrpA inhibits ComRS activity through disruption of ComR-XIP dimer formation, leading to measurable decreases in P*comX* activity, in ComX production and ultimately in a muted activation of competence (**Figure 9**).

**Figure 8.**
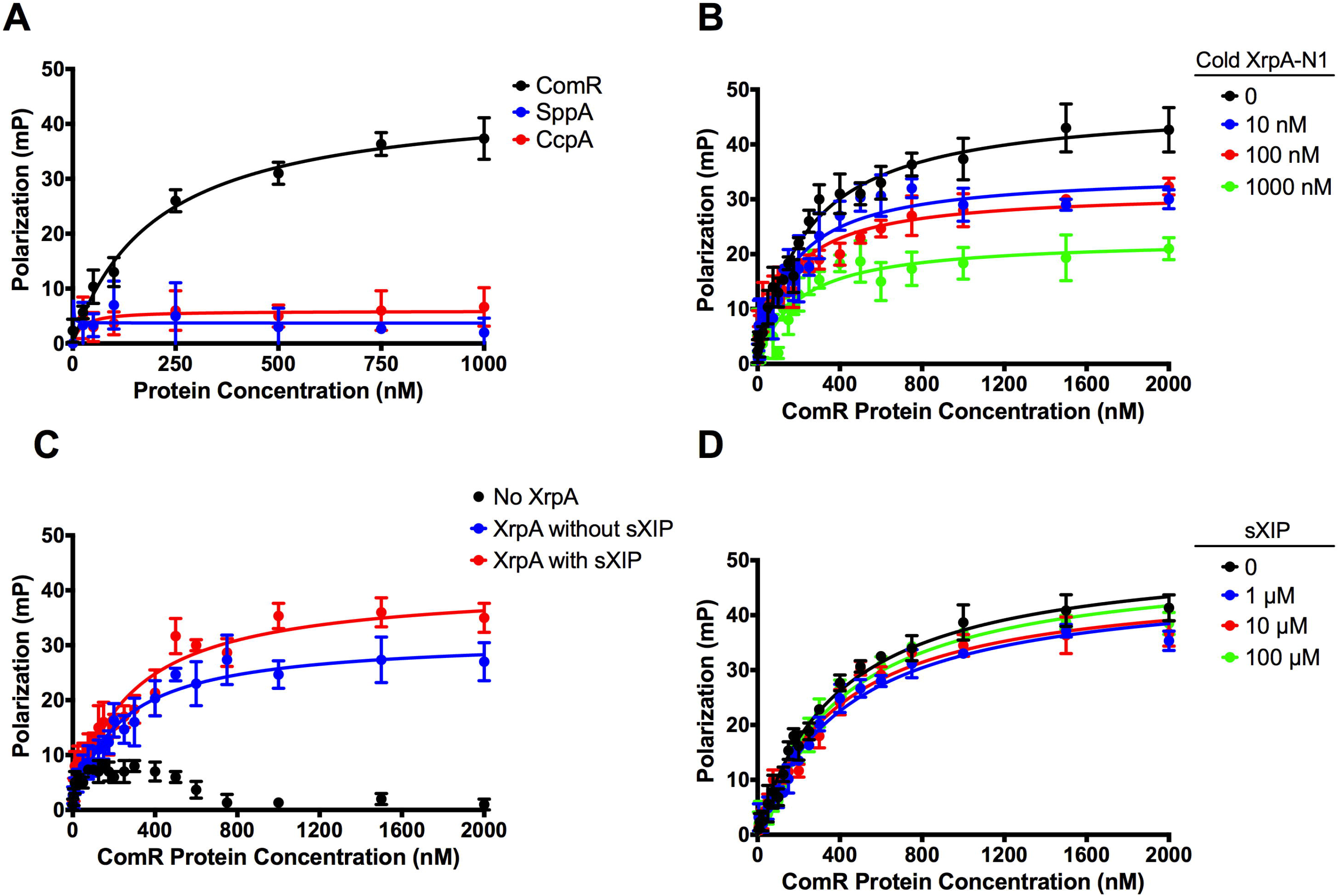
*Fluorescence Polarization experiments with labeled XrpA-N1*. (A) Selectivity of FITC-labeled XrpA-N1 probe to ComR (black) over other *S. mutans* purified recombinant proteins SppA (blue) and CcpA (red). Fluorescent polarization (FP) curves were generated using increasing concentrations of purified proteins binding to 10 nM of fluorescently labeled XrpA-N1 probe in the presence of 10 μM sXIP. (B) Increasing concentrations of synthetic unlabeled XrpA-N1 peptide were assessed for their ability to compete with FITC-labeled XrpA-N1 for their binding to increasing concentrations of purified ComR. (C). Binding of 10 nM FITC-labeled XrpA-N1 peptide to increasing concentrations of purified ComR, with and without 10 μM sXIP addition. (D) Binding of 10 nM FITC-labeled XrpA-N1 peptide to increasing concentrations of purified ComR in addition to increasing concentrations of sXIP peptide. Data shown for all graphs are averages of three independent experiments.

**Figure 9.**
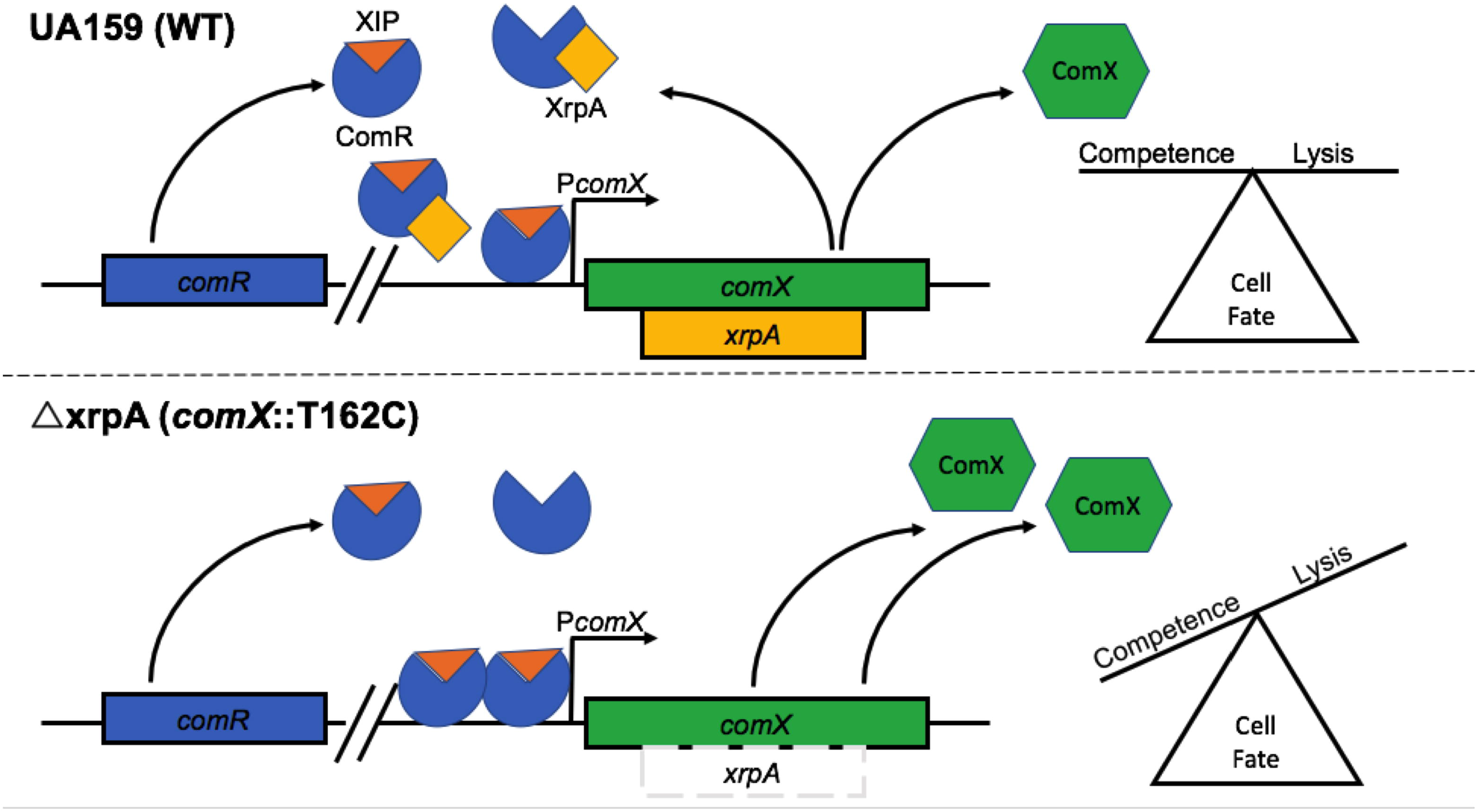
*Model for XrpA Modulation of ComRS Signaling*. Current model for the role of XrpA in inhibition of ComRS activity and in cell fate. When present and active, XrpA (yellow) interacts with ComR (blue) independently of whether ComR is in a complex with XIP (orange), resulting in diminished affinity of ComR for its target in PcomX. XrpA inhibition of ComR prevents the ComR-XIP complex from over-amplifying the competence activation signal, thereby maintaining a balance within the population of cells that induce competence and internalize DNA with a group of cells that undergo lysis providing DNA as a nutrient source, a source for genetic diversification, and/or a source of eDNA that contributes to extracellular matrix formation. One potential mechanism by which XrpA curtails proficient ComR-XIP activation of P*comX* is through inefficient dimer formation between ComR-XIP complexes. When XrpA is absent from the circuit, as in the case of the *xrpA* mutant (Δ*xrpA; comX*::T162C), over-amplification of ComRS signaling occurs. This leads to increases in the accumulation of the sigma factor ComX (green); which in turn results in an increase in the subpopulation of cells that undergo cell lysis.

## DISCUSSION

The study of bacterial cell-cell communication has provided valuable insights into bacterial processes that are critical for growth, essential for the activities of complex microbial ecosystems, and impactful of human health and diseases. These communication pathways have served as tractable model systems to dissect the intricacies of specialized secretory systems for signal molecules and bacteriocins, the mechanisms for signal transduction through two-component systems and cytosolic transcriptional regulators, the hierarchical control of regulons, and how multiple sensory inputs are integrated into key physiological outcomes and manifestation of virulence. As research has continued with these systems, the apparently straightforward paradigms once described for control of quorum sensing and intercellular communication have been assimilated into increasingly complex and diverse models for cellular reprogramming. Examples of such complexity can be found with negative feedback associated with the late competence gene *dprA* of *Streptococcus pneumoniae* (Weng *et al*., 2013), identification of the novel protein Kre that controls the bimodal regulation of ComK in *Bacillus subtilis* (Gamba *et al*., 2015), and the recent discovery of a short, leaderless, intercellular peptide signal in Group A *Streptococcus* (Do *et al*., 2017) that regulates protease expression. Similar advances in understanding of regulatory systems have been realized using *S. mutans* as a model organism (Lemos *et al*., 2013). One important characteristic that distinguishes *S. mutans* from other streptococci is that it encodes the ComRS signaling systems and integrates the ComCDE bacteriocin production pathway with competence through at least two different signal peptides XIP and CSP, respectively. Here we demonstrate additional complexities in the competence pathway of *S. mutans*, which is intimately intertwined with stress tolerance (Kaspar *et al*., 2016), by providing experimental evidence that a novel negative feedback system involving the unusual XrpA peptide is a regulator of the ComRS pathway and, consequently, of activation of late competence genes and lytic behaviors.

Using genetic and biochemical approaches, we confirmed that XrpA serves as a negative regulator of competence development in *S. mutans* by inhibiting activation of the targets of the ComR-XIP complex, apparently through a direct interaction with ComR that can be demonstrated *in vivo* by cross-linking and *in vitro* using purified constituents. We first described XrpA as an antagonist of ComX (Kaspar *et al*., 2015) that, based on its unusual genomic location within the *comX* gene and the inverse relationship of xrpA-specific mRNA abundance to the full-length *comX* transcript and ComX protein levels, might function as an anti-sigma factor, as opposed to acting on early competence genes (ComDE or ComRS systems). Instead, using transcriptome profiling and transcriptional reporter assays a picture began to emerge that XrpA influenced ComRS-dependent activation of the *comX* promoter. XrpA functioning as an autogenous negative regulator of its own expression by blocking ComR-dependent activation of *comX* may provide the cells with an opportunity to fine-tune ComS and ComX production, particularly in response to environmental inputs, such as redox. A similar type of regulation has been described for the *E. coli* RNA polymerase-binding protein DksA, which along with the co-factor (p)ppGpp promotes a negative feedback loop on the *dksA* promoter to keep DksA protein levels constant in different environmental conditions (Chandrangsu *et al*., 2011). Not only is it intriguing in evolutionary terms that the XrpA negative feedback system evolved within the *comX* coding region, but also there are potentially important physiological ramifications of the existence of this regulatory circuit. The exact mechanism by which the *xrpA* mRNA is translated has not been established, but we presently favor a model by which ribosomal slippage occurs during *comX* translation at or near the *xrpA* start codon, allowing for production of XrpA, with the efficiency of translational initiation at the *xrpA* start codon and the stability of the 5’ region of the *comX* mRNA being factors that govern the ratio of ComX to XrpA. Confirmation of such a model is the subject of ongoing research. Nevertheless, if we accept the premise that *xrpA* translation is not as efficient as that of *comX* when the *comX* promoter is activated, then it can be envisioned that the negative feedback loop created by XrpA acting on ComR modulates transcriptional initiation at the *comX* promoter; with the feedback loop contributing to fine tuning ComX levels in response to cellular physiology and environment and/or serving as a primary pathway to turn off the competence circuit. The former would be consistent with the observation that signal perception, induction of *comX* and progression to the competent state are all exquisitely sensitive to key environmental inputs that include pH (Guo *et al*., 2014; Son *et al*., 2015), oxidative stressors (De Furio *et al*., 2017) and carbohydrate source and availability (Moye *et al*., 2016). The latter model would be consistent with the fact that, while the MecA-Clp pathway can serve as a mechanism to shut off competence through degradation of ComX, the cells also need a way to shut down the ComRS circuit so as not to produce nascent ComX during competence while committing to turning off the competence regulon.

Our model for XrpA acting at the level of ComR-dependent activation of *comS* and *comX* is supported by the transcriptional reporter data in which overexpression of *xrpA* lead to decreased P*comX* activity (Figure 2). Recently it was suggested that an antagonist termed “ComZ” must be present within ComRS-positive streptococci that can shut down ComRS activity in a similar manner to DprA of the ComDE systems (Mirouze *et al*., 2013; Haustenne *et al*., 2015). We do not suspect that XrpA is the aforementioned ComZ, as the loss of *xrpA* only exhibits stronger activation of ComRS-dependent promoters, but does not increase the duration of activation, as is evident in Figure 2. Studies into ComX stability over the course of competence activation revealed that ComX protein accumulates faster at early time points after induction with signal peptides (10-30 minutes) in an xrpA-negative strain, consistent with the data on promoter activity, but ComX protein dissipates at a similar rate in the presence or absence of *xrpA* (data not shown). Thus, we conclude that XrpA modulates the strength of ComRS signaling, but other factors must be present, possibly working in concert with XrpA, to shut off competence in *S. mutans*, further highlighting the complexity of these systems. There is potential for other protein candidates identified in our SPINE experiment, such as *ssb2*, fulfill this role; an area that remains to be investigated.

In *S. mutans*, competence activation has been linked to cell lysis in a process similar to the fratricide that was first described for *S. pneumoniae*. Fratricide appears to be essential for efficient gene transfer between bacteria in biofilm communities and to be mediated in *S. mutans* through the activities of encoded cell wall hydrolases that are a part of the ComX regulon (Wei and Håvarstein, 2012; Khan *et al*., 2016), and possibly by intracellular bacteriocins (Perry *et al*., 2009). As seen in **Figure 3**, the inhibition of ComRS activity by XrpA has the ability to change the proportion of cells exhibiting responses to signal inputs in a population, both in terms of competence activation in complex medium and cell lysis, as measured by propidium iodide staining of XIP-treated cultures and production of eDNA, which in *S. mutans* is heavily dependent on cell lysis (Liao *et al*., 2014). We propose a model (Figure 9) in which *xrpA* influences the decision pathway by which cells choose between a) viability and potential to uptake DNA as a result of competence activation or b) fratricide and cell death, the latter providing a source of genetic material and eDNA for incorporation into the extracellular matrix during biofilm formation (Liao *et al*., 2014). We posit that this decision network relies on the strength and duration of P*comX* activation, although we cannot yet rule out the post-transcriptional regulatory factors (RNA binding proteins, RNAses, or riboswitching) govern the ratio of XrpA to ComX. Notwithstanding, when XrpA accumulates and/or is active, the strength of ComRS activation is moderated, providing an even balance between cell viability and cell death. In the absence of XrpA, ComRS-dependent activation causes ComX over-accumulation, which favors lytic behavior, which can be observed in Figure 3 and is supported by the fact that higher concentrations of exogenously supplied XIP induce cell death (Wenderska *et al*., 2012). Previously, we reported that overexpression of *xrpA* results in a growth defect in the presence of oxygen, linking *xrpA* expression and production to oxidative stress tolerance (Kaspar *et al*., 2015). One aspect not studied here is how *xrpA* might sense an oxidative environment and integrate that response into competence activation through modulation of ComRS activity. The C-terminal portion of XrpA, which does not appear to interact with ComR, is very hydrophobic and may be membrane-associate (Kaspar *et al*., 2015). Additionally, XrpA is unusual in that it contains seven cysteine residues distributed fairly evenly across the protein that could participate in disulfide bond formation, between XrpA proteins/peptides to influence XrpA availability or to form covalent interactions with binding partners. ComR contains three cysteine residues distributed evenly over its length, which could allow for covalent coupling to XrpA or sub-fragments of XrpA. It is our working hypothesis that XrpA integrates the oxidative state of the environment into fine-tuning the strength of the ComRS signal, tempering activation in one condition over the other. In the case of early biofilm formation, oxygen is readily available and could be sensed by environmental inputs such as XrpA, leading to its inactivation. In this scenario, high ComRS activation and ComX accumulation could shift a larger population of the cells into a lytic mode during competence activation, releasing eDNA to facilitate the formation of a protective extracellular matrix, thereby conveying increased fitness or persistence of *S. mutans* over other health-associated commensal streptococci. As oxygen levels and redox potential decrease with biofilm maturation, the need for strong competence activation could be diminished, shifting the cells into a more stable growth mode with a smaller proportion of cells undergoing lysis. It is noteworthy also that bacteriocins are among the most highly up-regulated genes when *S. mutans* is growing in air, compared with the transcriptome of anaerobically growing cells (Ahn *et al*., 2007). We are currently exploring these ideas and how the competence regulon is integrated with biofilm development and its role in competition with commensal streptococci. It is critical to note that ComX and XrpA are highly conserved in all sequenced clinical isolates of *S. mutans* (Kaspar *et al*., 2015), suggesting evolutionary pressure to keep these pathways intact. Our working hypothesis is this evolutionary pressure arises from the need to compete with commensal streptococci that can antagonize the growth of *S. mutans* through a variety of mechanisms (Bowen *et al*., 2017).

We have previously noted that *xrpA* appears to be unique to *S. mutans* (Kaspar *et al*., 2015). Indeed, a tblastn search for identification of ORFs with similar sequences bacteria showed that only *Streptococcus troglodytae*, a recently sequenced oral isolate from chimpanzees that is most closely related to *S. mutans* (Okamoto *et al*., 2013), contains an intact *xrpA* coding sequence embedded within the *comX* coding region. *Streptococcus dysgalactiae* subsp. *equisimilis* (SDSE) also contains a similar *xrpA* coding sequence, but the *xrpA* protein coding sequence is disrupted by premature stop codons in the sequenced isolate of this *Streptococcus*. Thus, it appears that only *S. mutans* and extremely closely related organisms are the only ComRS-containing streptococci that encode an XrpA-like inhibitor; there is no evidence that XrpA is present in *S. rattus, S. sobrinus. S. cricetus, S. downeii* or other mutans streptococci. We cannot, however, exclude that there are proteins or peptides in ComRS-containing streptococci that play a role that is similar or identical to that of XrpA in *S. mutans*. The unique nature of XrpA in *S. mutans* is also notable in the context that the *S. mutans* ComR has strict recognition for its cognate XIP peptide (Shanker *et al*., 2016), whereas ComR proteins from Bovis and Pyogenic streptococci are more promiscuous in the XIP peptides that they are able to recognize to enhance ComR DNA binding capacity. It is then a logical conclusion that the Mutans group has further separated from the Bovis and Pyogenic group in terms of ComR-XIP regulation. A logical extrapolation of these observations is to ask whether XrpA could serve as a novel anti-caries target or therapeutic. It is important to note that other oral health-associated commensal streptococci, such as *Streptococcus mitis, Streptococcus gordonii* and *Streptococcus sanguinis* are all part of the Mitis group of streptococci that lack ComRS signaling systems and rely on ComDE for competence activation (Håvarstein, 2010), so targeting XrpA should not disrupt a healthy oral biofilm. It is encouraging that small synthesized portions of XrpA have an effect on competence activation, as shown in Figure 4. It is critical to point out that while theses synthesized peptides were provided exogenously in this study, we do not suspect at this time that XrpA is actively released to the extracellular environment. As previously discussed, our current working hypothesis is that XrpA is able to accumulate in response to external environmental cues, such as oxidative stress, and function as a sensor inside the cells, and at this time have no reason to believe that XrpA has an extracellular lifecycle. Our working model must also account for the fact that XrpA fragments can elicit effects when provided exogenously. Therefore, we are testing the hypothesis that XrpA peptides, which may be released into the extracellular space through lysis, can be actively internalized, perhaps after processing, to elicit their effects on ComR – similar to what has been proposed for XIP (Kaspar *et al*., 2017). Mass spectrometry studies are currently ongoing to localize XrpA and other *S. mutans* encoded peptides that affect competence (Ahn *et al*., 2014).

In summary, this work provide additional novel insights into the complex regulatory nature of bacterial cell-cell signaling systems, providing the organisms with multiple check points throughout the circuit to either amplify or diminish the response to signal inputs based on key environmental and/or physiologic cues. Future work will be focused on how environmental inputs can influence XrpA/ComR interaction and activities, and the resulting consequences in terms of biofilm ecology. Development of these model systems should shed further light on microbial interactions and the importance of cell-cell signaling systems at the very earliest stages of colonization and biofilm development. It is also interesting to ponder if the recently discovered peptides such as XrpA, the rcrQ-associated peptides of *S. mutans* (Ahn *et al*., 2014) and the leaderless SHP of GAS (Do *et al*., 2017) play important roles in the virulence potential of these organisms if multiple other peptides with profound impacts on cellular behaviors are currently hidden in the genomes of Gram-positive bacteria.

## EXPERIMENTAL PROCEDURES

### Bacterial Strains and Growth Conditions

*S. mutans* wild-type strain UA159 and its derivatives (**Table 1**) were grown in either brain heart infusion (BHI - Difco), FMC (Terleckyj *et al*., 1975; Terleckyj and Shockman, 1975) or CDM (Chang *et al*., 2011) medium. The medium was supplemented with 10 μg ml^−1^ erythromycin, 1 mg ml^−1^ of kanamycin or 1 mg ml^−1^ spectinomycin when needed. Unless otherwise noted, cultures were grown overnight in BHI medium with the appropriate antibiotics at 37°C in a 5% CO_2_ aerobic atmosphere. The next day, cultures were harvested by centrifugation, washed twice in 1 mL of phosphate-buffered saline (PBS), and resuspended in PBS to remove all traces of BHI. Cells were then diluted in the desired medium before beginning each experiment. Synthetic XIP (sXIP, aa sequence = GLDWWSL), corresponding to residues 11-17 of ComS, was synthesized and purified to 96% homogeneity by NeoBioSci (Cambridge, MA). The lyophilized sXIP was reconstituted with 99.7% dimethyl sulfoxide (DMSO) to a final concentration of 2 mM and stored in 100 μL aliquots at −20°C. Selected XrpA peptide sequences and fluorescently-labeled derivatives were synthesized, purified and confirmed by mass spectrometry by Biomatik USA (Wilmington, DE). For scrambled peptides, the Shuffle Protein tool from www.bioinformatics.org was utilized. XrpA peptides were also reconstituted with DMSO to a final concentration of 1 mM and stored at −20°C.

**TABLE 1.** List of strains

| Strain or Plasmid | Relevant Characteristics | Source or Reference |
| --- | --- | --- |
| *S. mutans* Strains |  |  |
| UA159 | Wild-type | ATCC 700610 |
| $\Delta xrpA$ | Inactivation of *xrpA* start codon;<br>*comX*::T162C | (Kaspar *et al.*, 2015) |
| 184XrpA/UA159 | UA159 harboring overexpression of *xrpA* on pIB184, Em<sup>R</sup> | (Kaspar *et al.*, 2015) |
| $\Delta comR$ | *comR* (SMU.61) :: Em<sup>R</sup> | (Kaspar *et al.*, 2016) |
| $\Delta comX$ | *comX* (SMU.1997) :: Em<sup>R</sup> | (Kaspar *et al.*, 2015) |
| PcomX::gfp/UA159 | UA159 harboring *gfp* fluorescent reporter of PcomX | (Son *et al.*, 2012) |
| PcomX::gfp/$\Delta xrpA$ | $\Delta xrpA$ harboring *gfp* fluorescent reporter of PcomX | This study |
| PcomX::gfp/184XrpA | 184XrpA/UA159 harboring *gfp* fluorescent reporter of PcomX | This study |
| PcomS::gfp/UA159 | UA159 harboring *gfp* fluorescent reporter of PcomS | (Kaspar *et al.*, 2017) |
| PcomS::gfp/$\Delta xrpA$ | $\Delta xrpA$ harboring *gfp* fluorescent reporter of PcomS | This study |
| PcomS::gfp/184XrpA | 184XrpA/UA159 harboring *gfp* fluorescent reporter of PcomS | This study |
| PcomX::gfp/$\Delta comS$ | $\Delta comS$ (*comS* :: Em<sup>R</sup>) harboring *gfp* fluorescent reporter of PcomX | (Son *et al.*, 2012) |
| PcomX::gfp/$\Delta comS\Delta xrpA$ | $\Delta comS$ in *comX*::T162C harboring *gfp* fluorescent reporter of PcomX | This study |
| 184ComR-Strep/UA159 | UA159 harboring overexpression of *comR* with addition of (G4S)<sub>2</sub> linker and Strep-tag on pIB184, Em<sup>R</sup> | This study |
| Plasmids |  |  |
| pDL278 | Escherichia coli - Streptococcus shuttle vector, Sp<sup>R</sup> | (LeBlanc *et al.*, 1992) |
| pIB184 | Shuttle expression plasmid with the constitutive P23 promoter, Em<sup>R</sup> | (Biswas *et al.*, 2008) |

\* Em, erythromycin; Sp, spectinomycin.

**TABLE 2.**
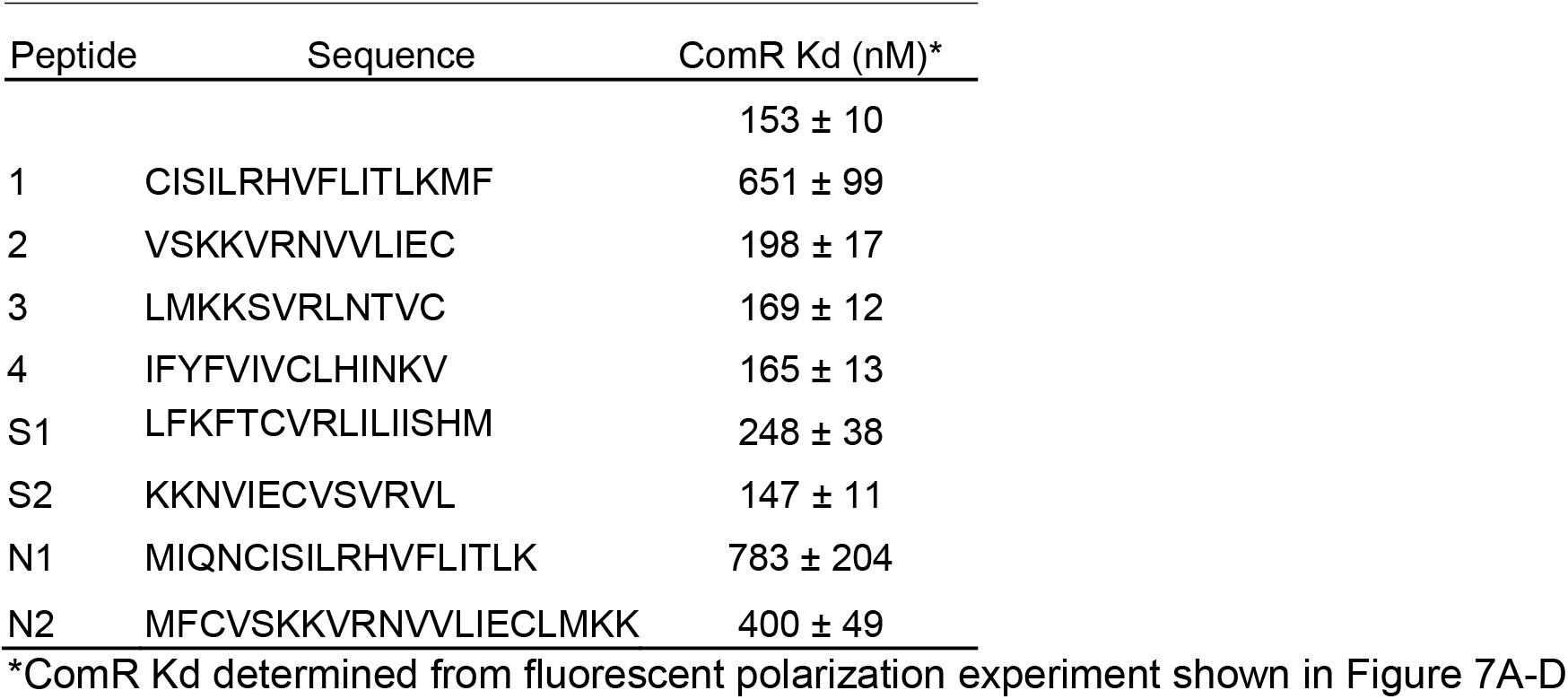
ComR Kd from Fluorescent Polarization

### Construction of Bacterial Strains

Mutant strains of *S. mutans*, including inactivation of the *xrpA* start codon (Δ*xrpA*) were created using a PCR ligation mutagenesis approach as previously described (Lau *et al*., 2002; Kaspar *et al*., 2015). Overexpression of genes (*xrpA, comR*) was achieved by amplifying the structural genes of interest from *S. mutans* UA159 and cloning into the expression plasmid pIB184 using the EcoRI and BamHI restriction sites (Biswas *et al*., 2008). For *in vivo* protein-protein interaction experiments, a Strep-tag sequencing (WSHPQFEK) was first inserted in front of the stop codon on the pIB184-ComR overexpressing plasmid using the Q5^®^ Site Directed Mutagenesis Kit (New England Biolabs, Beverly, Mass.) and following the provided protocol. After selection of the appropriate construct by sequencing, a [G4S]_2_ Linker sequence (ggtggaggaggctctggtggaggcggtagc) was then inserted between the *comR* and Strep-tag sequence using the same kit and protocol (**Supplemental Table 1**). Transformants were confirmed by PCR and sequencing after selection on BHI agar with appropriate antibiotics. Plasmid DNA was isolated from *E. coli* using QIAGEN (Chatsworth, Calif.) miniprep columns, and restriction and DNA-modifying enzymes were obtained from New England Biolabs. PCRs were carried out with 100 ng of chromosomal DNA by using *Taq* DNA polymerase, and PCR products were purified with the QIAquick kit (QIAGEN).

### Transcriptome Profiling via RNA-Seq

Selected strains of *S. mutans* to be analyzed by RNA-sequencing (UA159, Δ*xrpA*) were grown in FMC medium to mid-exponential log phase of OD_600_ = 0.5 before harvesting. For *S. mutans* UA159 treated with 2 μM sXIP, sXIP was added at OD_600_ = 0.2. RNA extraction, rRNA removal, library construction and read analysis was conducted as previously described elsewhere (Zeng *et al*., 2013; Kaspar *et al*., 2015). Briefly, 10 μg of high-quality total RNA was processed using the MICROB*Express*™ Bacterial mRNA Enrichment Kit (Ambion of Life Technologies, Grand Island, NY), twice, before ethanol precipitation and resuspension in 25 μL of nuclease-free water. The quality of enriched mRNA samples was analyzed using an Agilent Bioanalyzer (Agilent Technologies, Santa Clara, CA). cDNA libraries were generated from the enriched mRNA samples using the TruSeq Illumina kit (Illumina, San Diego, CA), following instructions from the supplier. Deep sequencing was performed at the University of Florida ICBR facilities (Gainesville, FL). Approximately 20 million short-reads were obtained for each sample. After removing adapter sequences from each short-read and trimming of the 3’-ends by quality scores (Schmieder and Edwards, 2011), the resulting sequences were mapped onto the reference genome of strain UA159 (GenBank accession no. AE014133) using the short-read aligner. Mapped short-read alignments were then converted into readable formats using SAMTOOLS (Li *et al*., 2009). For viewing of the mapped reads aligned to the genome, .bam files were uploaded into the Integrative Genomics Viewer (IGV – version 2.3.55) (Robinson *et al*., 2011). A “.csv” file containing raw read counts for each replicate (3) was then uploaded to Degust (http://degust.erc.monash.edu/) and edgeR analysis performed to determine Log2 fold change and a false discovery rate (FDR). The P-value was obtained by taking the −log10 of the FDR. The data files used in this study are available from NCBI-GEO (Gene Expression Omnibus) under accession no. GSE110167.

### Measurements of Promoter Activity via GFP Fluorescence

For measurements of GFP fluorescence, cultures were inoculated from washed overnight cultures in CDM medium at a 1:50 dilution. Inoculated medium (175 μL) was added to each well along with a 50 μL mineral oil overlay in a Costar™ 96 well assay plate (black plate with clear bottom; Corning Incorporated) and incubated at 37°C. At intervals of 30 minutes for a total of 18 hours, OD_600_ along with GFP fluorescence (excitation 485/20 nm, emission 528/20 nm) was measured with a Synergy 2 multimode microplate reader (BioTek). Relative expression was calculated by subtracting the background fluorescence of UA159 (mean from six replicates) from raw fluorescence units of the reporter strains and then dividing by OD_600_.

### Flow cytometry

Bacterial cultures were grown to OD_600_ = 0.6 before being harvested, washed and resuspended in PBS before being run through a FACSCalibur™ (BD Biosciences) flow cytometer. For sCSP experiments, cultures were grown in BHI after a 1:20 dilution from overnight culture while cultures were grown in FMC for sXIP experiments at the same initial dilution. Both peptides were added to the growing cultures at OD_600_ = 0.2. sXIP treated cells were stained with 5 μg mL^−1^ propidium iodide (PI) for 10 minutes in the dark for analysis of membrane-compromised cells. Cells were then sonicated in a water bath sonicator for 3 intervals of 30 seconds in 5 mL polystyrene round-bottom tubes to achieve primarily single cells for analysis. Forward and side scatter signals were set stringently to allow sorting of single cells. In total, 5 × 10^4^ cells were counted from each event, at a maximum rate of 2 × 10^3^ cells per second, and each experiment was performed in triplicate. Detection of GFP fluorescence was through a 530 nm (± 30 nm) bandpass filter, and PI was detected using a 670-nm long pass filter. Data were acquired for unstained cells and single-color positive controls so that data collection parameters could be properly set. The data were collected using Cell Quest Pro (BD Biosciences) and analyzed with FCS Express 4 (De Novo Software). Gating for quadrant analysis was selected by using a dot density plot with forward and side scatter, with gates set to capture the densest section of the plot. Graphing and statistical analyses were performed using Prism (GraphPad Software). x- and y-axis data represent logarithmic scales of fluorescent intensity (arbitrary units).

### Measurements of eDNA Release

Overnight cultures of selected *S. mutans* strains, grown in CDM medium with addition of a final concentration of 10 μM of synthetic XrpA peptides when noted, where measured for a final OD_600_ and then harvested by centrifugation for their supernatant fraction. 5 mL of the resulting supernatant was then run through QIAGEN (Chatsworth, Calif.) PCR purification columns to capture eDNA present. The eDNA was then eluted off the column with 600 μL water, and 594 μL of this elution was mixed with 5 μL of 50 μM Sytox Green (Invitrogen) to a final concentration of 0.5 μM. After vortexing the solution and incubation for 15 minutes in the dark at room temperature, 200 μL of the stained samples were transferred into a Costar™ 96 well assay plate (black plate with clear bottom; Corning Incorporated). Fluorescence (excitation 485/20 nm, emission 528/20 nm) was measured with a Synergy 2 multimode microplate reader (BioTek) and the resulting data was then normalized for the measured final OD _600_ nm resulting in a final arbitrary eDNA release measurement. The data represents 3 independent biological replicates with 3 technical replicates each. Statistical significance was determined by the Student’s T-Test.

### Biofilm Assays

Selected *S. mutans* strains were grown from overnight cultures to mid-exponential phase after a 1:20 dilution in BHI broth at 37°C in a 5% CO_2_ atmosphere. The mid-exponential phase cells were then diluted 1:100 into CDM medium with containing 15 mM glucose and 2.5 mM sucrose as a carbohydrate source. 200 μL of this dilution was loaded into 96 well polystyrene microtiter plates and incubated in a 5% CO_2_ atmosphere at 37°C for 48 h. After, the medium was decanted, and the plates were washed twice with 200 μL of sterile water to remove planktonic and loosely bound cells. The adherent bacteria were stained with 60 μL of 0.1% crystal violet for 15 min. After rinsing twice with 200 μL of water, the bound dye was extracted from the stained biofilm using 200 μL of ethanol:acetone (8:2) solution, twice. The extracted dye was diluted into 1.6 mL of ethanol:acetone solution. Biofilm formation was quantified by measuring the absorbance of the solution at OD _575_ nm. The data represents 3 independent biological replicates with 4 technical replicates each. Statistical significance was determined by the Student’s T-Test.

### SPINE for ComR Interactions

A Strep-tag protein interaction experiment (SPINE) was derived from a previously published protocol (Herzberg *et al*., 2007). Briefly, a strain harboring a C-terminal Strep-tagged ComR, along with the vector only control, was grown in 500 mL of CDM medium after a 1:20 dilution from overnight cultures to an OD600 = 0.6. At this time, either 2 μM of sXIP or DMSO (vehicle, 0.1% final concentration) was added to cultures and were grown an additional hour at 37°C in a 5% CO_2_ atmosphere. After, cells were pelleted by centrifugation, washed and resuspened in 50 mM HEPES (pH 8) with a final concentration of 2.5 mM of protein crosslinker solution added (DSP, Thermo Scientific). The cells were incubated at 37°C for 45 minutes, at which point 50 mM Tris (pH 7.5) was added to stop the crosslinking reaction. After, cells were pelleted, washed in Buffer W, and lysed via bead beating in Buffer W. The crosslinked Strep-tagged ComR complex was then purified from the lysate using Strep-tactin^®^ resin (iba) in a chromatography column following the manufacture’s protocol. Purification was verified by running the selected fractions and elutions on 16.5% Tris-Tricine gels (BioRad) followed by silver staining and/or western blot (**Supplemental Figure 1**). Protein concentrations of the elutions were determined using the bicinchoninic acid assay (BCA; Thermo Scientific). Complex-containing elutions were then combined and precipitated by the TCA/Acetone precipitation method. Precipitant was sent to Applied Biomics (Hayward, CA) for 2D DIGE Protein Expression Profiling which included 2D gel electrophoresis, determination of protein expression ratios between samples, spot picking and identification by LC-MS/MS though trypsin digestion of selected spots then returned peptide fragments compared to a list of predicted masses using a *S. mutans* UA159 database of all annotated proteins.

### Transformation Assays

Overnight cultures were diluted 1:20 in 200 μL of FMC medium in polystyrene microtiter plates in the presence or absence of 10 μM of synthetic XrpA peptides. The cells were grown to OD_600_ = 0.15 in a 5% CO_2_ atmosphere. When desired, 0.5 μM of sXIP was added, cells were incubated for 10 min and 0.5 μg of purified plasmid pIB184, which harbors a erythromycin resistance (Erm^R^) gene, was added to the culture. After 2.5 h incubation at 37°C, transformants and total CFU were enumerated by plating appropriate dilutions on BHI agar plates with and without the addition of 1 mg mL^−1^ erythromycin, respectively. CFU were counted after 48 h of incubation, and transformation efficiency was expressed as the percentage of transformants among the total viable cells. Fold change was then calculated from the UA159 control with DMSO (vehicle) addition. Statistical significance was determined by the Student’s T-Test

### Western blot

Overnight cultures of *S. mutans* were diluted 1:50 into 35 mL of FMC medium and harvested by centrifugation when the cultures reached an OD_600_ = 0.5. When desired, 2 μM sXIP was added when the cultures reached an OD_600_ value of 0.2 along with 10 μM of selected XrpA peptides. Cell pellets collected by centrifugation were washed once with buffer A (0.5 M sucrose; 10 mM Tris-HCl, pH 6.8; 10 mM MgSO4) containing 10 μg mL^−1^ of phenylmethanesulfoynl fluoride (PMSF) (ICN Biomedicals Inc.) and resuspended in 0.5 mL Tris-buffered saline (50 mM Tris-HCl, pH 7.5; 150 mM NaCl). Cells were lysed using a Mini Bead Beater (Biospec Products) in the presence of 1 volume of glass beads (avg. diam. 0.1 mm) for 30 s intervals, three times, with incubation on ice between homogenizations. Lysates were then centrifuged at 3,000 × *g* for 10 minutes at 4°C. Protein concentrations of the resulting supernates were determined using the bicinchoninic acid assay (BCA; Thermo Scientific) with purified bovine serum albumin as the standard. Ten microgram aliquots of proteins were mixed with 5X SDS sample buffer (200 mM Tris-HCl, pH 6.8; 10% [v/v] SDS; 20% [v/v]; 10% [v/v] β-merchaptoethanol; 0.02% [v/v] bromophenol blue), loaded on a 12.5% polyacrylamide gel with a 5% stacking gel and separated by electrophoresis at 150 V for 45 minutes. Proteins were transferred to Immobilon-P polyvinylidene difluroride (PVDF) membranes (Millipore) using a Trans-Blot Turbo transfer system and a protocol provided by the supplier (BioRad). The membranes were treated with either primary polyclonal anti-ComX, anti-ComR or anti-ManL (loading control) antisera at a 1:1000 dilution and a secondary peroxidase-labeled, goat anti-rabbit IgG antibody (1:5000 dilution; Kierkegaard & Perry Laboratories, USA). Detection was performed using a SuperSignal West Pico Chemiluminescent Substrate kit (Thermo Scientific) and visualized with a FluorChem 8900 imaging system (Alpha Innotech, USA).

### Purification of Recombinant ComR

The *S. mutans* UA159 *comR* gene was amplified using primers AAAGAATCCTATGTTAAAAGA and CACCCTAGGAGACCCATCAAA and was cloned into the BamHI and AvrII sites of the pET 45b expression vector downstream of the 6x His-tag and separated by an enterokinase cleavage site. The resulting vector was transformed into *E. coli* DE3 cells (New England Biolabs). To induce expression of *comR*, 1 mM isopropyl-β-D-thiogalactopyranoside (IPTG) was added to 1 L of growing culture in LB medium once the OD_600_ reached 0.6. The culture was grown for an additional 4 h at 37°C before the cells were pelleted by centrifugation and frozen overnight at −20°C. The next day, the cells were lysed after suspension into B-PER (Thermo Scientific) with addition of HALT protease inhibitor (Thermo Scientific) and cellular debris removed by centrifugation for 20 minutes at 13,000 x g. The His-ComR was column purified using Ni-NTA resin (QIAGEN) and eluted with 250 mM imidazole (Supplemental Figure 4). To remove the 6x His-tag, 1 mL of a 2-3 mg mL^−1^ purified His-ComR sample was dialyzed in EKMax Buffer overnight (50 mM Tris-HCL pH 8, 1 mM CaCl_2_, 0.1% Tween-20) with 50 U of EKMax enterokinase (Thermo Scientific) then added and incubated overnight at 4°C. Finally, the cleaved ComR sample was added to a dialysis cassette to exchange the EKMax buffer with PBS pH 7.4 overnight. Final protein concentration was determined using the bicinchoninic acid assay (BCA; Thermo Scientific) with purified bovine serum albumin as the standard. Digestion was confirmed by SDS-PAGE and Coomassie Blue staining (Supplemental Figure 5).

### Fluorescence Polarization

A 5’ Bodipy-labeled self-annealing, stem-loop DNA probe with sequence encompassing the ECom-box which ComR binds to within the P*comX* promoter (5’-BODIPY FL-X - ATGGGACATTTATGTCCTGTCCCCCACAGGACATAAATGTCCCAT - 3’), was synthesized (Thermo Fisher) and kept at a constant concentration of 1 μM in all reactions. Purified ComR protein was serially diluted, ranging from 5 to 2000 nM, and mixed with 10 μM sXIP and 10 μM of selected synthetic XrpA peptides unless otherwise noted, in reaction buffer to a final volume of 250 μL (PBS pH 7.4, 10 mM βME, 1 mM EDTA, 0.1 mg mL^−1^ BSA, 20% glycerol (v/v), 0.01% Triton X-100 and 0.05 mg mL^−1^ salmon sperm DNA). Reactions were transferred to a Corning^®^ 96-well, half-area, black polystyrene plate prior to incubation at 37°C for 30 minutes.

Polarization values were measured using a BioTek Synergy 2 plate reader (excitation 485 nm, emission 528 nm), and the resulting millipolarization values were plotted for each protein concentration tested to assess protein/peptide interactions. For fluorescent peptides experiments, a synthetic fluorescein isothiocyanate (FITC)-labeled XrpA-N1 peptide (FITC-AHX-MIQNCISILRHFLITLK) was used (Biomatik USA) with reaction buffer PBS pH 7.4, 10 mM βME, 0.1 mg mL^−1^ BSA, 20% glycerol (v/v), and 0.01% Triton X-100. Graphing and linear regression analyses to determine kD values were performed using Prism (GraphPad Software).

## ACKNOWLEDGEMENTS

Research reported in this publication was supported by the National Institute of Dental and Craniofacial Research of the National Institutes of Health under Award Numbers R01 DE13239, R01 DE023339, and T90 DE21990.

## AUTHOR CONTRIBUTIONS

JK, RCS and RAB contributed to the conception and design of the study; JK and RCS performed the experiments, acquired and analyzed the data, JK and RAB interpreted the data; and JK and RAB wrote the manuscript.

